# Ongoing habenular activity is driven by forebrain networks and modulated by olfactory stimuli

**DOI:** 10.1101/2021.02.14.431141

**Authors:** Ewelina Magdalena Bartoszek, Suresh Kumar Jetti, Khac Thanh Phong Chau, Emre Yaksi

## Abstract

Ongoing neural activity, which represents internal brain states, is constantly modulated by the sensory information that is generated by the environment. In this study, we show that the habenular circuits act as a major brain hub integrating the structured ongoing activity of the limbic forebrain circuitry and the olfactory information. We demonstrate that ancestral homologs of amygdala and hippocampus in zebrafish forebrain are the major drivers of ongoing habenular activity. We also reveal that odor stimuli can modulate the activity of specific habenular neurons that are driven by this forebrain circuitry. Our results highlight a major role for the olfactory system in regulating the ongoing activity of the habenula and the forebrain, thereby altering brain’s internal states.

## INTRODUCTION

Continuous interactions between the sensory world and the internal states of the brain are essential for survival. For example, a sudden change in the environment by the presence of a salient sensory cue can be critical for updating an animals’ brain from its resting to an alert state [1–3]. Seminal studies in human subjects clearly demonstrated the presence of such rapid alterations from a default state with high levels of brain activity [4, 5] to an alert state with reduced levels of global network activity and increased activity in sensory or attention related areas during various tasks [3, 6, 7]. While the precise definition of a ‘brain state’ can vary depending on the time scales of investigated phenomena, the link between ongoing (or spontaneous) brain activity and the current state of the brain is well accepted [8–14]. In fact, ongoing neural activity has been observed across the nervous system, from brain areas involved in sensory processing [15–23], to innate behaviors [24–26], and higher cognitive function [27–37]. Despite the presence of ongoing activity across the brain and its rapid alterations coupled with brain state transitions, dedicated neural pathways and their connection to ongoing brain activity and sensory-evoked transitions remain to be identified.

One brain region that receives information from both sensory [38–40] and cortico-limbic [41–54] brain structures, and exhibits high level of ongoing neural activity, is the habenula. This evolutionary conserved diencephalic nucleus [44, 55–57] is a major hub relaying information from diverse forebrain inputs [41, 43–48, 50, 58] to its target regions that directly control animal behavior via the release of dopamine, acetylcholine, and serotonin [40, 44, 45, 59–69]. Previous studies established a direct role for the habenula in adaptive behaviors [8, 64, 65, 70–74], learning [45, 47, 64, 73, 75, 76] and prediction of outcomes [45, 71, 75, 77, 78]. Not surprisingly, habenular dysfunction is closely linked to several neurological conditions and mood disorders including depression [54, 79]. Based on molecular profiles [80–85], anatomical landmarks [64, 73, 76, 86], and neural activity [52, 53, 64, 87], habenula is divided into numerous subdomains. Most prominent in zebrafish are the dorsal (dHb) and the ventral habenula (vHb), which are the homologous to mammalian lateral and medial habenula [64, 73, 76, 86], respectively. dHb is involved in sensory processing [40, 67, 88], circadian rhythms [67], social behaviors [72] and experience-dependent fear response [65, 73, 76]. Whereas vHb plays important roles in learning [64] and active coping behavior [8]. Recent studies showed that ongoing neural activity associated with the transition between brain states is present in both dHb and vHb [8, 52, 53, 65]. Interestingly, sensory cues that have been shown to induce prominent behavioral responses in zebrafish [8, 65, 66, 83, 89–91], were also shown to elicit distinct neural responses both in dHb [39, 52, 65, 87] and vHb [8, 52]. Yet, it is less clear how ongoing habenular activity is generated by the distributed brain networks and how this ongoing activity interacts with sensory responses in the habenula and across the brain.

In this study, we investigated how interactions between ongoing and sensory-evoked activity can shape brain’s internal states, in the habenula and across the entire zebrafish forebrain. First, we showed that functional interactions of neurons within distinct habenular subdomains are stable and spatially organized across different periods of ongoing activity. Next, we revealed that ongoing habenular activity is driven by the ancestral homologs of limbic forebrain regions, hippocampus and amygdala. Finally, we demonstrated that olfactory cues switch the internal states of the habenula by specifically modulating the activity of habenular neurons driven by ancestral limbic forebrain regions. Our results reveal that the habenula acts as a hub integrating limbic and sensory signals, and olfactory cues switch internal brain states by inhibiting the ongoing activity of the habenula and its ancestral limbic forebrain inputs.

## RESULTS

### Structured ongoing activity in the habenula is stable over time

Toinvestigateongoinghabenularactivity,weperformed volumetric two-photon calcium imaging across the entire habenula of juvenile Tg(elavl3:GCaMP6s) zebrafish [52, 92–94], expressing GCaMP6s panneuronally. In all our experiments, we used 3 to 4-weeks-old zebrafish that were shown to exhibit complex behaviors such as learning [75, 95, 96] and social interactions [97, 98], which are mediated by habenular circuits [64, 72, 73, 75, 78]. We observed structured ongoing activity across the entire habenula (Figure 1A), in line with earlier studies focusing on the dHb [53]. To quantify ongoing habenular activity, we performed k-means clustering on habenular calcium signals and observed that distinct functional clusters of habenular neurons exhibit synchronous activity (Figure 1A, B, Figure S1). Next, we investigated, whether such functional clusters in habenula are stable across different time periods, by comparing the activity of functional clusters that were recorded in two consecutive time windows. We quantified the stability of habenular clusters by measuring the probability of a pair of habenular neurons in the same cluster during period #1 to remain in the same clusters in period #2, which we termed “cluster fidelity” [52, 53]. We observed that more than 40% of habenular neuron pairs remained in the same cluster, which is significantly above chance levels (Figure 1C). The functional clusters of habenular neurons identified by k-means clustering appeared to be spatially organized across the entire habenula (Figure 1B, Figure S1). To quantify the spatial distribution of the synchronous habenular activity, we plot the average pairwise correlation of neurons versus the distance between them. We observed that nearby habenular neurons exhibit more correlated/synchronous ongoing activity, when compared to distant pairs of neurons within each habenular hemisphere (Figure 1D). Our results using k-means clustering of ongoing habenular activity suggest that while neurons within the same clusters exhibit highly correlated activity (i.e. within individual clusters of Figure 1A), neurons in different clusters can also be anti-correlated (i.e. between cluster #3 and cluster#5 of Figure 1A). We investigated whether such correlations and anti-correlations between habenular neurons are stable across different time periods, by plotting pairwise correlations of all recorded neuron pairs. In fact, we observed that correlations and anti-correlations between habenular neurons remained stable across different time periods (Figure 1E, black), and this relationship was different from shuffled distributions of such pairwise relationships (Figure 1E, grey). Taken together, these results revealed that the ongoing habenular activity is highly structured, stable over time, and spatially organized into functional clusters of habenular neurons exhibiting synchronous or anti-synchronous ongoing activity.

**Figure 1.**
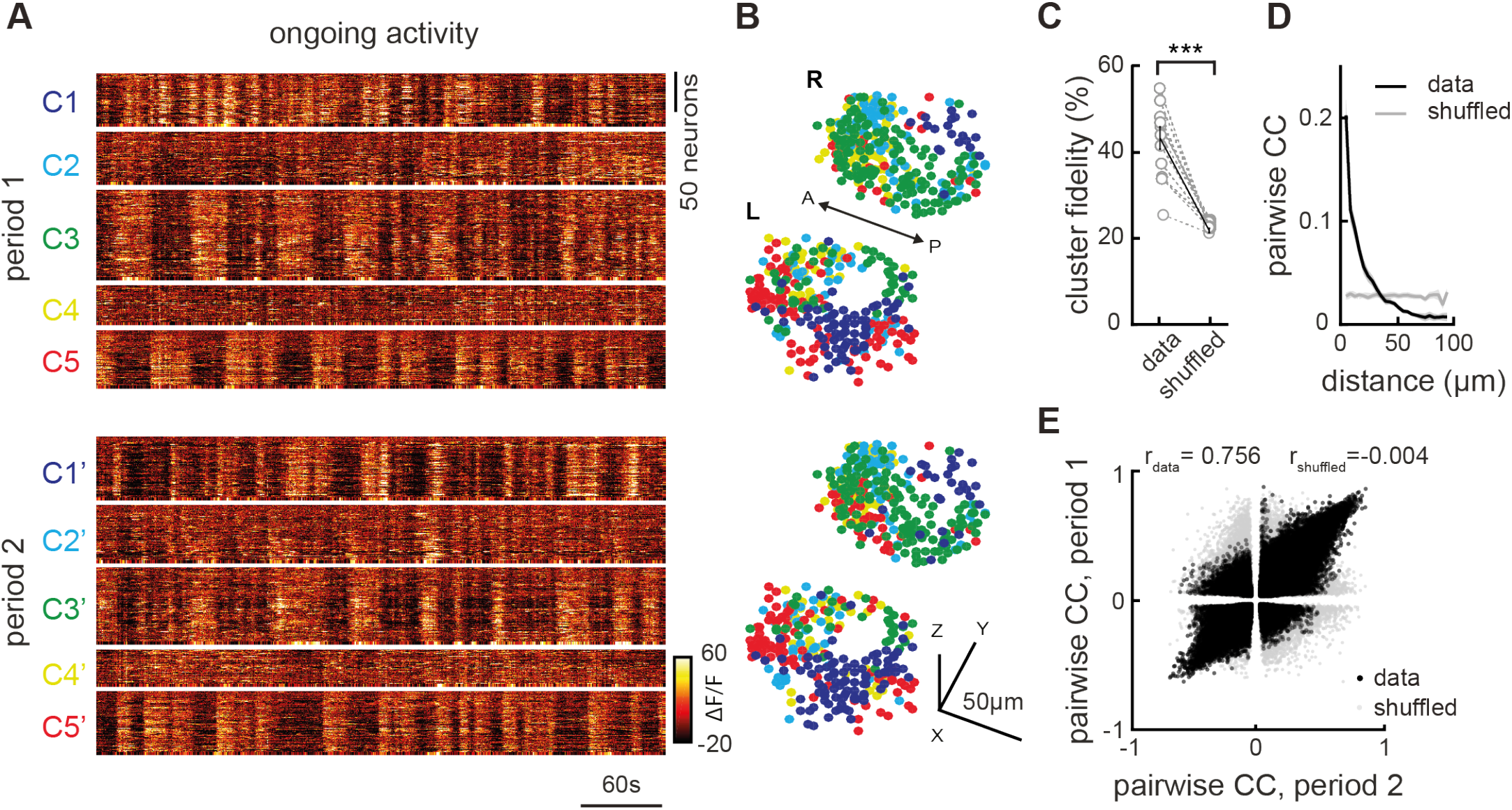
Ongoing activity of habenular neurons is temporally and spatially organized. (A) Representative example of ongoing habenular activity recorded by two-photon calcium imaging in Tg(eval3:GCaMP6s) zebrafish line. Habenular neurons are clustered (C1-5) using k-means clustering over two consecutive time periods of 7 minutes each (period 1 – top and period 2 – bottom). Warm colors represent higher calcium signals. (B) Representative example of three-dimensional reconstruction of habenular neurons clustered using k-means clustering in two consecutive time periods (top, bottom). Neurons are color-coded based on their cluster identity that correspond to the calcium signals depicted in A. L – left; R – right hemisphere, A – anterior, P – posterior. (C) The ratio of habenular neuron pairs remaining in the same functional clusters (high cluster fidelity) is significantly higher than chance levels, during two different time periods of ongoing activity. 315 ± 35 (mean ± SEM) habenular neurons were imaged in each fish (n=11 fish). ***p<0.001, Wilcoxon signed rank test. (D) Relation between pairwise correlation of habenular neurons during ongoing activity and the distance between each neuron pair. Grey line represents shuffled spatial distributions. (E) Pairwise correlations of calcium traces of habenular neurons (with p-value<0.05) during two consecutive time periods, in black. Grey dots represent pairwise comparison that are shuffled for pair identities. Actual data exhibit a correlation of rdata=0.756 for the pairwise correlation across two time periods, indicating robust synchrony between pairs of neurons. Shuffled distribution is rs=−0.004.

### Distinct forebrain regions correlate with ongoing habenular activity

Previous studies in mammals [45, 47, 48, 50, 99–103] and in zebrafish [39, 41, 52, 53, 65, 67, 72, 73, 75, 80, 87] showed that several forebrain regions send anatomical projections to the habenula. We asked which of these candidate forebrain regions might drive and modulate ongoing habenular activity. To do this, we measured the ongoing activity of the entire forebrain of the juvenile zebrafish, including the habenula, by using volumetric two-photon calcium imaging. To identify forebrain regions that might drive distinct functional clusters of habenula, we asked which forebrain neurons are most correlated with the average ongoing activity of individual habenular clusters (Figure 2A, B). We observed that the neurons of dorsolateral (Dl), dorsomedial (Dm), ventrodorsal forebrain (Vd), and the olfactory bulbs (OB) contained neurons with highest correlations to individual habenular clusters (Figure 2C, D, Figure S2).

**Figure 2:**
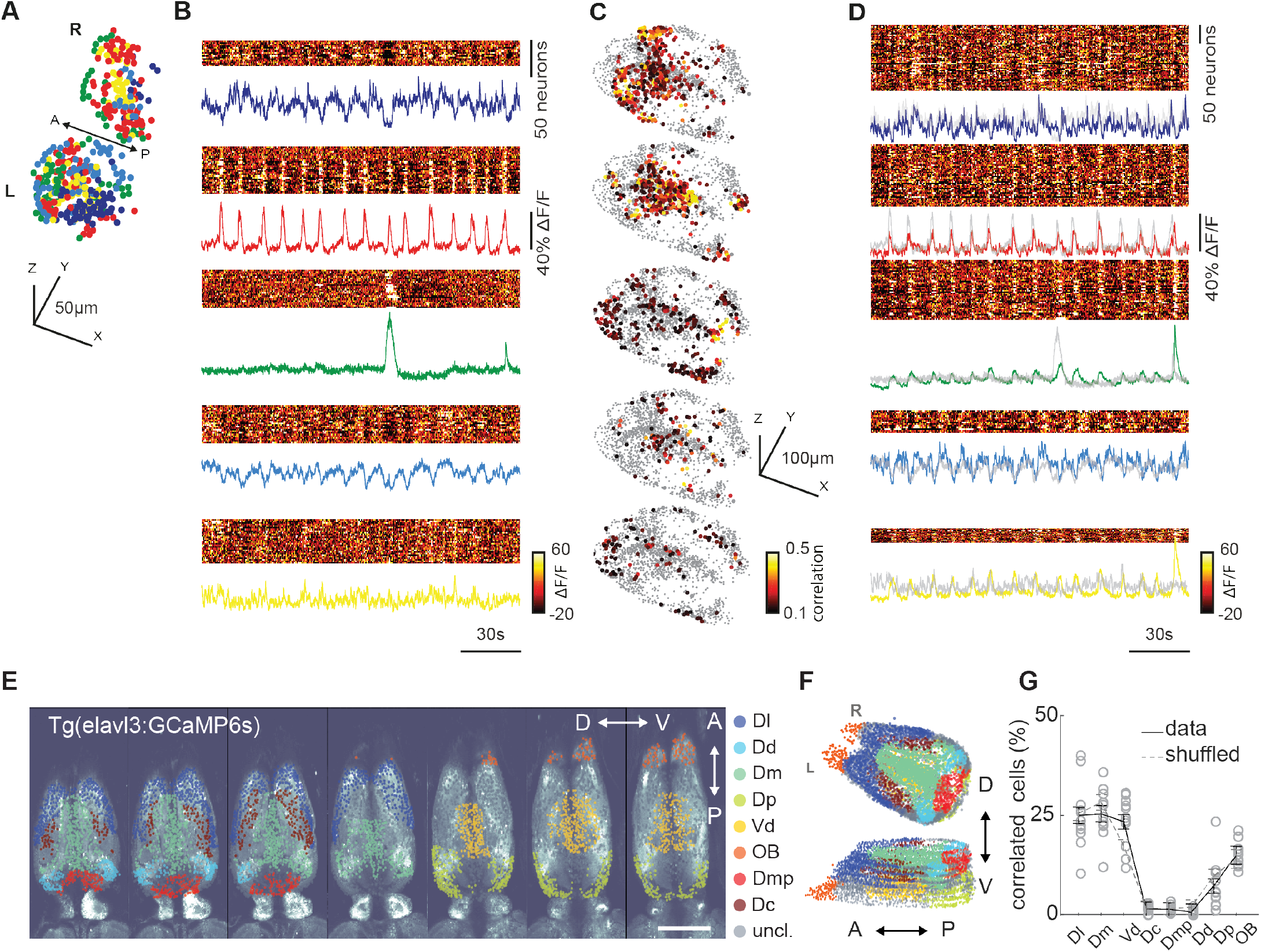
Ongoing activity of habenular neurons is correlated with sensory and limbic forebrain regions. (A) Reconstruction of habenular neurons in Tg(elavl3:GCaMP6s) zebrafish, clustered with k-means clustering. Colors represent clusters with similar ongoing activity. L-left; R-right hemisphere. 315 ± 35 (mean ± SEM) habenular neurons were imaged in each fish (n=11). (B) Ongoing activity of the habenular neurons corresponding to clusters in A. Warm colors represent higher activity. Color-coded traces represent the average activity of all neurons in each cluster. (C) Reconstruction of forebrain neurons that are strongly correlated (Pearson’s correlation >0.1) to average ongoing activity of habenular clusters in B. Warm colors represent stronger correlations. 2135 ± 345 (mean ± SEM) forebrain neurons were imaged in each fish (n=11 fish). (D) Ongoing activity of the forebrain neurons corresponding to clusters of neurons depicted in C. Color-coded traces represent the average activity of neurons in each cluster. Grey traces represent the average activity of habenular clusters in B. Note that the ongoing activity of identified forebrain neurons and habenular clusters are highly similar. (E) Forebrain regions identified based on anatomical land marks are color coded, and overlaid on raw two-photon microscopy image. Scale bar represents 100um. Optical planes are shown from dorsal to ventral. A – anterior, P – posterior, D – dorsal, V – ventral. (F) Reconstruction of forebrain regions shown in E. (G) Distribution of forebrain neurons with strong correlation (>0.1) to ongoing habenular activity into anatomically identified forebrain regions. Dl – dorsolateral telencephalon, Dd – dorsal nucleus of the dorsal telencephalon, Dm – dorsomedial telencephalon, Dp – posterior zone of the dorsal telencephalon, Vd – dorsal nucleus of the ventral telencephalon, OB – olfactory bulb, Dmp – posterior nucleus of dorsomedial telencephalon, Dc – central zone of the dorsal telencephalon. Black lines represent mean ± SEM.

To identify and quantify the forebrain regions driving habenula, we manually delineated distinct forebrain nuclei using anatomical landmarks that were identified by previous studies in zebrafish [38, 104–113] and in other teleosts [114–122] (Figure 2E, F). To further test whether anatomically identified forebrain regions overlapped with functionally distinct forebrain clusters, we compared those manually delineated forebrain nuclei (Figure S3A, B) with k-means functional clusters of ongoing forebrain activity (Figure S3C). We quantified this overlap with a previously used cluster selectivity index [52, 53]. Cluster selectivity would be “0” if all neurons of an anatomically delineated nuclei were randomly distributed to different functional clusters, or “1” if all neurons of the nuclei were in the same functional cluster. In fact, our results showed that cluster selectivity is significantly higher than the chance levels for all forebrain regions (Figure S3D), suggesting that the neurons of manually delineated forebrain regions are selective to one or few functional clusters. Next, we asked in which of these forebrain regions we can find neurons that are strongly correlated (Pearson’s correlation above 0.1) with the average ongoing activity of each habenular cluster. Our results revealed that the dorsolateral (Dl), dorsomedial (Dm), and ventrodorsal (Vd) telencephalic regions, homologous to mammalian hippocampus [120, 121, 123–126], amygdala [106, 116, 124, 125, 127, 128] and striatum [55, 124, 129] respectively, contains the largest fraction of habenula correlated neurons, in addition to the olfactory pathway (Figure 2G). These results suggest that ancestral homologs of mammalian limbic forebrain circuits in zebrafish, Dl, Dm, Vd, are potential candidates to drive ongoing habenular activity.

### Electrical micro-stimulation of limbic forebrain regions Dl and Dm elicits spatially organized responses in the habenula

Our correlation-based functional connectivity measurements (Figure 2) suggest that distinct limbic forebrain regions might be the major drivers of ongoing habenular activity. To test this hypothesis directly, we locally activated dorsal forebrain by a glass micro-stimulation electrode while measuring forebrain and habenular activity, in a juvenile zebrafish brain-explant preparation with intact connectivity [93]. Inserting the micro-stimulation electrode in Dl or Dm allowed direct activation of these dorsal forebrain regions (Figure 3A, J). Micro-stimulation of Dl and Dm also elicit significant activation in 23-29% of habenular neurons (Figure 3B, C, K, L), that are spatially organized (Figure 3D, M). Application of glutamate receptor blockers (AP5/NBQX) silenced habenular responses (Figure S4), confirming that the habenular responses upon forebrain micro-stimulation is not due to direct electrical activation of habenula, but due to glutamatergic inputs received by the habenular neurons. Due to its intact connectivity, juvenile zebrafish brain-explant exhibits sufficient level of structured ongoing habenular activity to identify functional clusters in habenula by using k-means clustering (Figure 3E, F, N, O). We observed that ~40% of habenular neurons (significantly higher than chance levels) within a given functional cluster remained in the same cluster during forebrain micro-stimulation of Dm and Dl (Figure 3G-I, P-R). These results revealed the causal relationship between stimulation of forebrain regions Dm and Dl with the activation of habenular neurons, thereby confirming functional connections from these ancestral limbic dorsal forebrain structures onto the habenula. Our results also demonstrated that structured ongoing activity of habenula carries information from its input regions in the forebrain.

**Figure 3:**
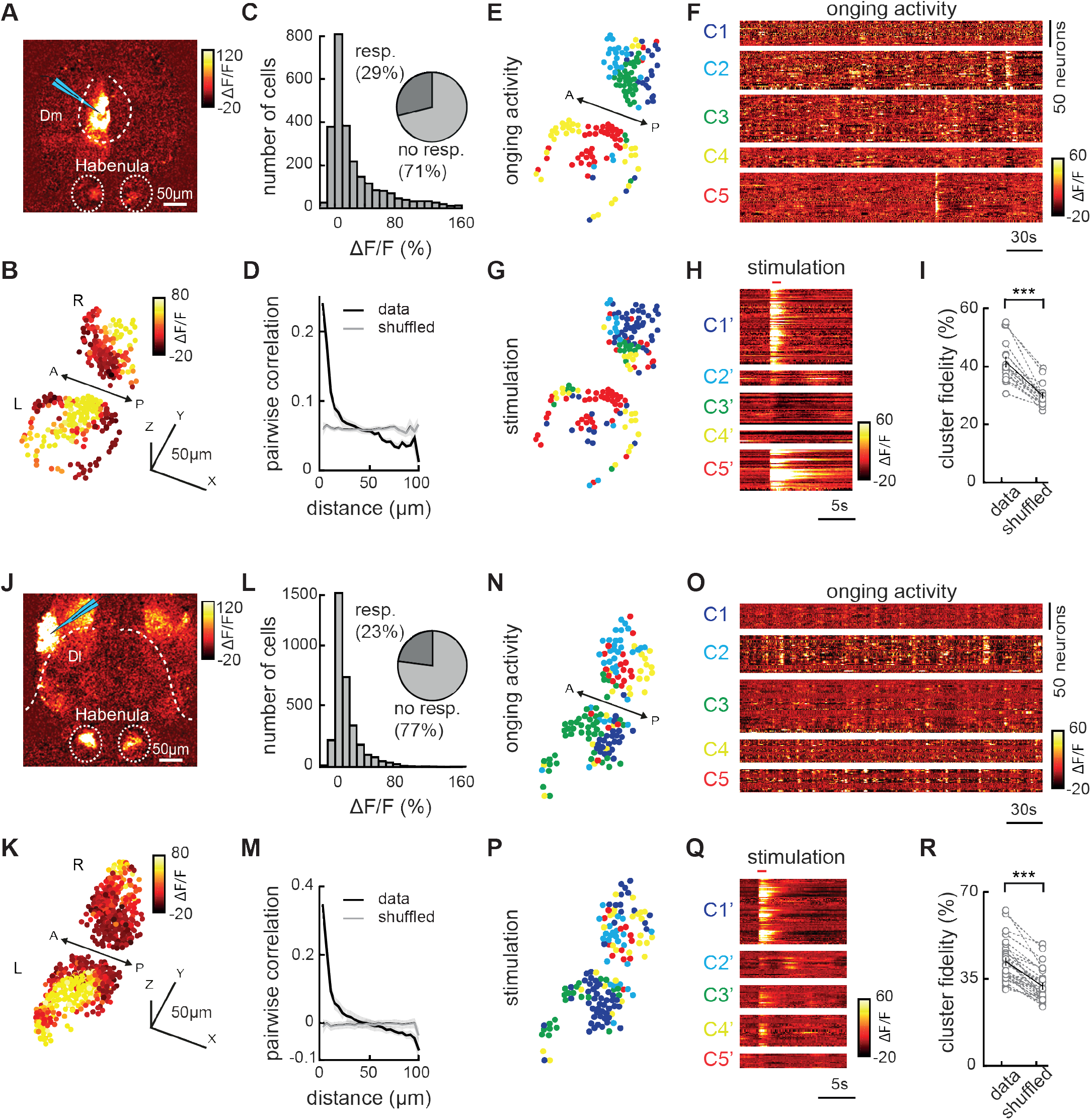
Electrical micro-stimulation of forebrain regions Dm and Dl activates spatially organized clusters of habenular neurons. (A) Two-photon calcium signals upon electrical micro-stimulation of forebrain region Dm, in Tg(eval3:GCaMP5) juvenile zebrafish brain explant. Warm colors represent stronger neural activity. Cyan triangle represents electrode location. Habenula and Dm are delineated with dashed lines. (B) Reconstruction of habenular responses upon Dm micro-stimulation. (C) Histogram representing the responses of habenular neurons upon Dm micro-stimulation (511±72 neurons per fish, in n=5 fish). Pie-chart represents the ratio of habenular neurons that are activated 2 SDs higher than baseline levels, upon Dm micro-stimulation. (D) Relation between pairwise correlation of habenular neuron responses upon Dm micro-stimulation and the distance between each neuron pair. Grey line represents shuffled spatial distribution. (E) Reconstruction of habenular neurons clustered with k-means functional clustering of their ongoing activity in juvenile zebrafish brain explant. Colors represent habenular clusters with similar ongoing activity. Scale bar represents 50μm. L – left; R – right hemisphere. (F) Ongoing activity of the habenular neurons corresponding to clusters in E. (G) Reconstruction of habenular neurons clustered during Dm micro-stimulation using k-means clustering. Colors represent habenular clusters with similar responses to Dm micro-stimulation. L – left; R – right hemisphere. (H) Responses of habenular neurons upon Dm micro-stimulation clustered by k-means clustering. Forebrain micro-stimulations is marked in red. Warm colors represent higher neural activity. (I) The ratio of habenular neuron pairs remaining in the same functional clusters (high cluster fidelity) is significantly higher than chance levels, during ongoing activity and Dm micro-stimulation. ***p<0.001, Wilcoxon signed rank test. (J) Two-photon calcium signals upon electrical micro-stimulation of forebrain region Dl, in Tg(eval3:GCaMP5) juvenile zebrafish brain-explant.. Cyan triangle represents electrode location. Habenula and Dl are delineated with dashed lines. (K) Reconstruction of habenular responses upon Dl micro-stimulation. (L) Histogram representing the responses of habenular neurons upon Dl micro-stimulation (575±92 neurons per fish, in n=6 fish). Pie-chart represents the ratio of habenular neurons that are activated 2 SDs higher then baseline levels upon Dl micro-stimulation. (M) Relation between pairwise correlation of habenular neuron responses upon Dl micro-stimulation and the distance between each neuron pair. Grey line represents shuffled spatial distribution. (N) Reconstruction of habenular neurons clustered with k-means functional clustering of their ongoing activity in juvenile zebrafish brain explant. Colors represent habenular clusters with similar ongoing activity. L-left; R-right hemisphere. (O) Ongoing activity of the habenular neurons corresponding to clusters in N. (P) Reconstruction of habenular neurons clustered during Dl micro-stimulation using k-means clustering. Colors represent habenular clusters with similar responses to Dl micro-stimulation. L – left; R – right hemisphere. (Q) Responses of habenular neurons upon Dl micro-stimulation clustered by k-means clustering. Forebrain micro-stimulations is marked in red. (R) The ratio of habenular neuron pairs remaining in the same functional clusters (high cluster fidelity) is significantly higher than chance levels, during ongoing activity and Dl micro-stimulation. ***p<0.001, Wilcoxon signed rank test.signed rank test.

### Sensory information and limbic dorsal forebrain inputs are integrated in the habenula

Previous studies showed that habenular neurons receive direct inputs from the olfactory bulbs [38, 111] and exhibit prominent chemosensory responses to diverse odors [39, 53, 87, 130]. Our functional connectivity analysis (Figure 2) and forebrain micro-stimulation experiments (Figure 3) also showed that at least a part of the ongoing habenular activity is generated by the activity of the olfactory pathway and the limbic forebrain regions Dm and Dl. We asked how these chemosensory and non-sensory forebrain inputs are integrated at the level of habenula. To do this, we established a nose-attached intact brain-explant preparation [131, 132] of juvenile zebrafish, which allowed micro-electrode stimulation of limbic dorsal forebrain regions, while delivering odors to the nose (Figure 4A). We observed habenular responses both upon odor delivery and dorsal forebrain micro-electrode stimulation (Figure 4B, C). We tested whether the habenula has distinct populations of neurons with different preferences for dorsal forebrain and olfactory inputs. Interestingly, a majority of the habenular neurons that are strongly activated by the micro-stimulation of the dorsal forebrain showed weak or no odor responses (Figure 4E), supporting the idea that habenula is composed of distinct zones with sensory or limbic inputs. Next, we asked whether chemosensory and limbic inputs to habenula can influence each other, when they are delivered simultaneously (Figure 4D). We first visualized the influence of dorsal forebrain stimulation on the habenular odor responses, by plotting the odor responses of habenular neurons against the change of their odor responses in the presence of forebrain stimulation (Figure 4F). In fact, 18.3% of odor responding habenular neurons showed a significant change in their odor responses upon forebrain stimulation (Figure 4F, black). The majority of these significantly modulated odor responses were weakly odor responding habenular neurons, whose responses were potentiated by forebrain stimulation. Next, we asked whether odor stimuli can modulate the responses of habenular neurons to forebrain stimulation. We observed that odor delivery significantly modulates forebrain driven responses in 21.7% of habenular neurons (Figure 4G, black), with a broad preference for inhibition of forebrain responses in the habenula, during odor stimulation (negative values in y axis, Figure 4G). These results indicate that the habenula integrates chemosensory and limbic information, and that these two different types of inputs can modulate each other, with a prominent suppression of limbic inputs during odor stimulation.

**Figure 4:**
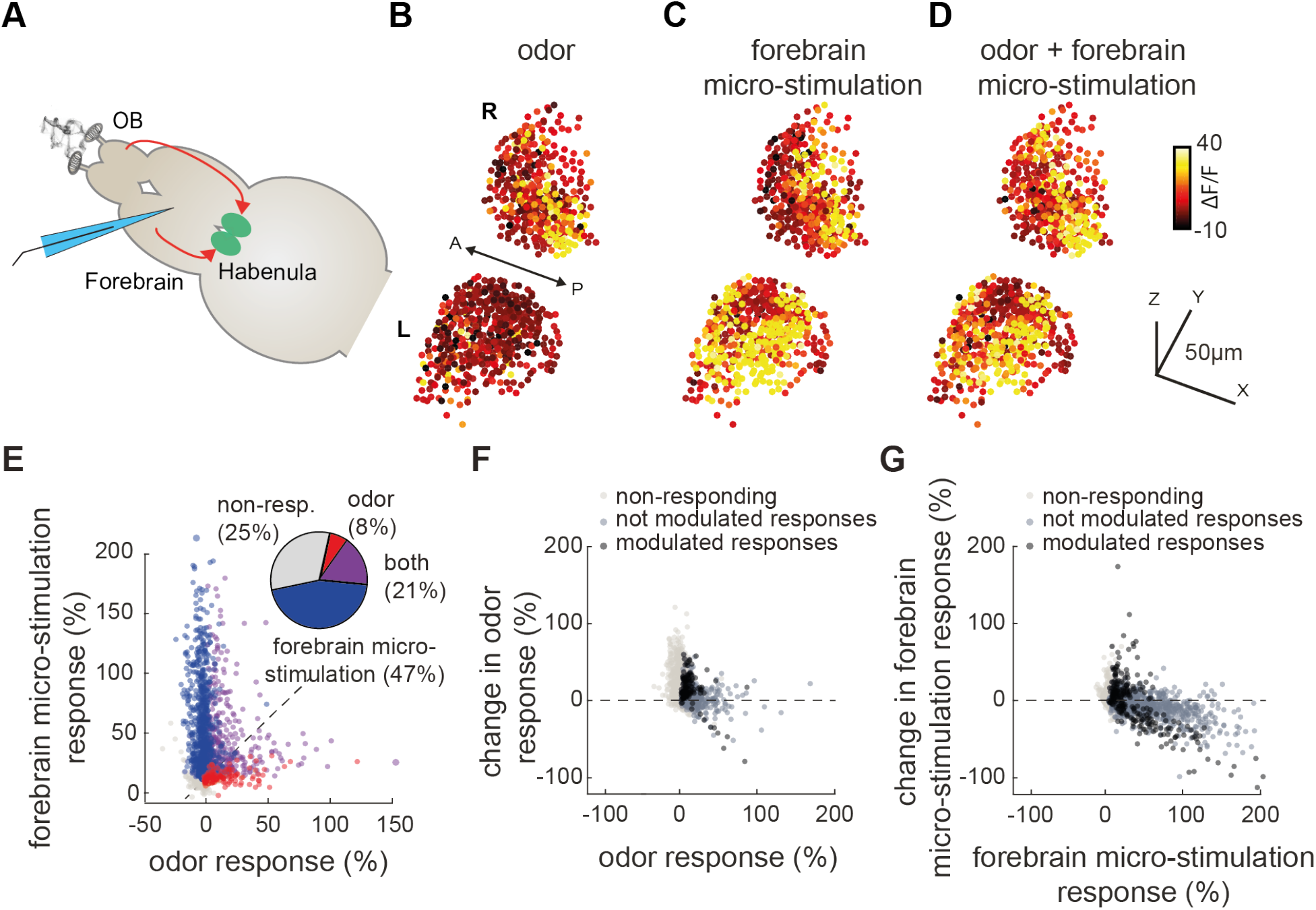
Habenular neurons integrate inputs from dorsal forebrain and olfactory system in a non-linear manner. (A) Schematic representation of nose-attached brain-explant preparation in Tg(elavl3:GCaMP6s) juvenile zebrafish, allowing simultaneous micro-stimulation of dorsal forebrain and odor stimulation of the nose. Habenula is marked in green, stimulation electrode is marked in cyan. Red arrows represent olfactory and dorsal forebrain inputs. (B) Three-dimensional reconstruction of habenular responses to odor stimulation averaged over 6 trials Warm colors indicate stronger neural responses. L – left; R – right hemisphere. (C) Reconstruction of habenular responses to dorsal forebrain micro-stimulation averaged over 6 trials. (D) Three-dimensional reconstruction of habenular responses to simultaneous odor stimulation and dorsal forebrain micro-stimulation averaged over 6 trials. (E) Responses of individual habenular neurons to odor stimulation and dorsal forebrain micro-stimulation from 2550 neurons measured in n=5 fish. Pie chart represents the ratio of habenular neurons responding 2 SDs above baseline levels to only odors (red), only micro-stimulation (blue) and both (magenta). (F) Change of odor responses in habenular neurons upon dorsal forebrain stimulation. Dark grey marks habenular neurons responding to odor stimulation. Black marks habenular neurons responding to odor stimulation and that are significantly (p<0.05, Wilcoxon signed rank test) modulated by forebrain micro-stimulation. (G) Change of habenular neuron responses to dorsal forebrain activation upon odor stimulation. Dark grey marks habenular neurons responding to dorsal forebrain micro-stimulation. Black marks habenular neurons responding to dorsal forebrain micro-stimulation and that are significantly (p<0.05, Wilcoxon signed rank test) modulated by the presentation of odors.

### Chemosensory stimulation modulates the ongoing activity of habenular neurons that are functionally coupled with limbic forebrain regions

Our results combining odor delivery with forebrain micro-stimulation in zebrafish brain-explant showed that chemosensory stimulation can modulate the activity of habenular neurons that receive forebrain inputs. We asked whether such odor-induced modulation of limbic forebrain-driven habenular neurons is present in vivo. To test this, we first identified the habenular neurons that are strongly (top 5%) inhibited (in blue) or excited (in red) by a range of attractive and aversive odors (Figure 5A, B), and calculated the average ongoing activity of these neurons (Figure 5C). Next, we identified the top 5% of forebrain neurons (Figure 5D), which are strongly correlated with the ongoing activity of habenular neurons that are either strongly inhibited or excited by odors. Finally, we determined how these forebrain neurons with strong correlations to odor modulated habenular neurons are distributed into distinct forebrain regions identified manually, as described in Figure 2F. Our results revealed that zebrafish homolog of hippocampus Dl, contains the largest number of neurons that are highly correlated with odor modulated habenular neurons (Figure 5E). In fact, Dl is the only brain region that contains neurons correlated with odor modulated habenular neurons (Figure 5E, blue and red labels) significantly more than the shuffled chance levels dictated by the size of individual regions (Figure 5E, dashed line). We did not observe any difference in the effect of aversive or attractive odors (Figure S5A, B). Our observations were also robust at different thresholds for selecting forebrain neurons strongly correlated with odor-modulated habenular neurons (Figure S5C, D).

**Figure 5:**
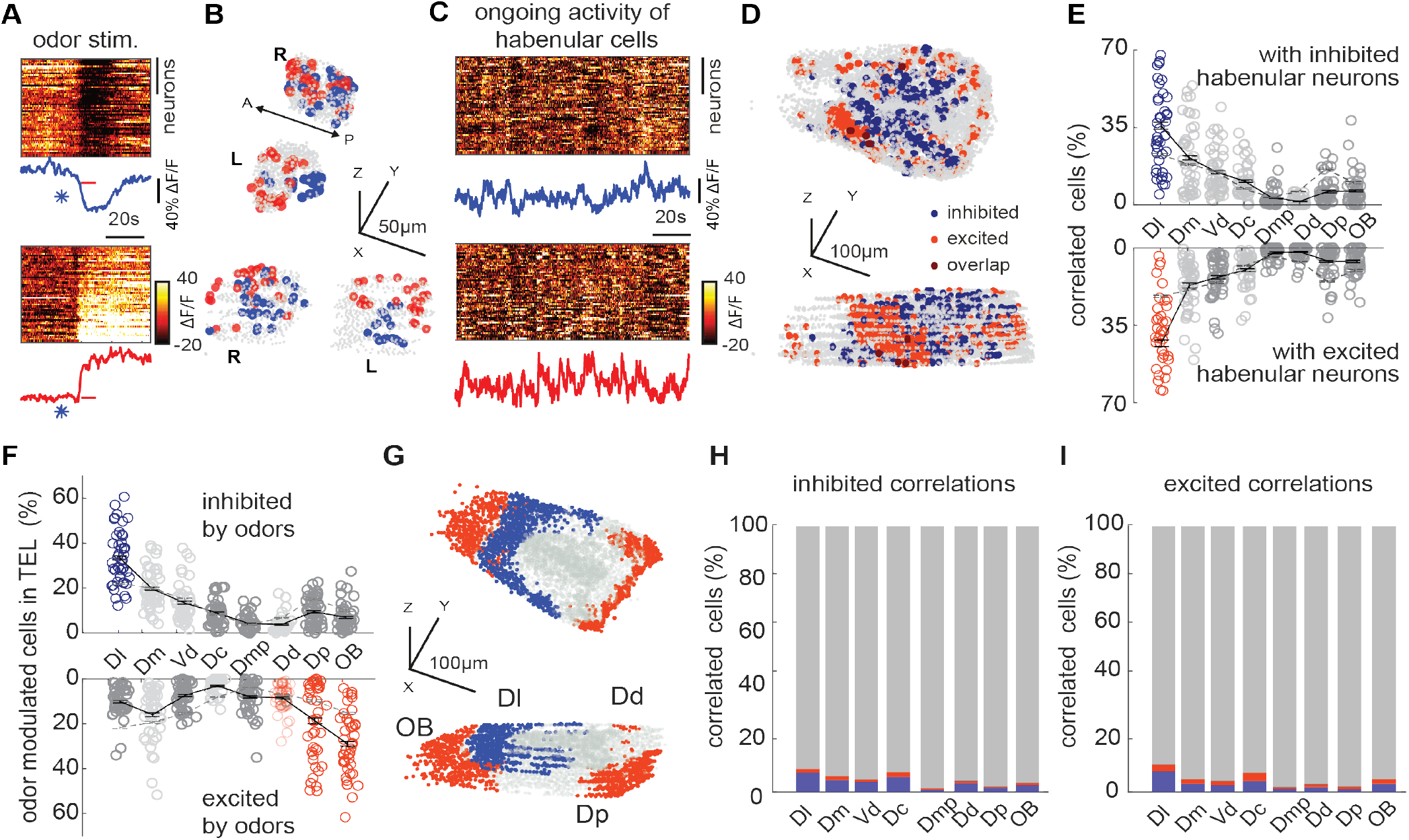
Odor-modulated habenular neurons are functionally connected with dorsal forebrain during ongoing activity. (A) Representative example of odor induced inhibition (top) and excitation (bottom) in habenular neurons of juvenile zebrafish. Dark colors represent inhibition (top 5% most inhibited neurons). Warm colors represent excitation (top 5% most excited neurons). Individual traces represent average inhibition (blue) and excitation (red) of habenular neurons. “*” indicates opening of odor valve, red bar indicates odor presentation. (B) Representative example of top 5% odor inhibited (blue) and excited (red) habenular neurons from an individual fish. 916 ± 39 (mean ± SEM) habenular neurons were imaged in each fish (n=10). (C) Ongoing neural activity of odor inhibited (top) and odor excited (bottom) habenular neurons in B. Lines represents the average activity of odor-inhibited (blue) and odor-excited (red) habenular neurons. (D) Representative example of top 5% forebrain neurons that are correlated with the ongoing activity of odor-inhibited (blue) and odor-excited (red) habenular neurons. 5799 ± 165 (mean ± SEM) forebrain neurons were imaged in each fish (n=10). (E) Anatomical distribution of forebrain neurons with strong correlations (top 5%) to ongoing activity of habenular neurons that are inhibited (top) and excited (bottom) by odors. Red and blue colored circles highlight forebrain regions that exhibit significantly higher synchrony to habenula than the chance levels dictated by the size of individual regions, p < 0.01, Wilcoxon signed-rank test. Black lines represent mean ± SEM. (F) Anatomical distribution of forebrain neurons that are most (top 5%) inhibited (top) and most excited (bottom) by odors. Colored circles highlight those forebrain regions that exhibit significantly higher number of odor-inhibited (blue) and odor-excited (red) neurons than the shuffled chance levels dictated by the size of individual regions, p < 0.01, Wilcoxon signed-rank test. Black lines represent mean ± SEM. (G) Representation of forebrain regions that show significantly higher number of odor-inhibited (blue) and odor-excited (red) neurons than the chance levels dictated by the size of individual regions. (H-I) Overlap of odor responding (blue: inhibited, red: excited) forebrain neurons and those forebrain neurons driving odor-inhibited (H) and odor-excited (I) habenular neurons.

Our results demonstrate the presence of strong chemosensory modulation (excitation/inhibition) of limbic-driven habenular activity. Yet, it is not clear if such chemosensory modulation happens at the level of habenular neurons or also at the level of limbic forebrain regions of zebrafish. To investigate the distribution of chemosensory modulation of dorsal forebrain neurons, we determined odor responses across the forebrain. Our results showed that Dl is the only forebrain region with odor-inhibited neurons significantly above chance levels (Figure 5F, top). As expected, olfactory bulb and olfactory cortex (Dp) [133], have the largest fraction of odor excited neurons, that are also significantly above the chance levels, in addition to dorsal zone of dorsal telencephalon (Dd) (Figure 5F, bottom). We observed no significant differences in the distribution of attractive or aversive odor responses (Figure S5C, D). The locations of forebrain regions that are strongly (top 5%) inhibited (blue) or excited (red) by odors are highlighted in Figure 5G.

Next, we asked how much of this odor-induced excitation and inhibition across the forebrain can explain the odor modulation of habenular activity. We observed that only a small fraction (less than 10%) of odor-excited or - inhibited forebrain neurons, overlap with forebrain neurons that drive odor-modulated habenular neurons (Figure 5H, I). This finding suggests that odors can separately modulate the limbic forebrain and habenular neurons with limited functional connections. Altogether, our results revealed that odors inhibit the activity of the ancestral limbic forebrain circuits in zebrafish. We also showed that odor-induced modulation of habenular activity cannot solely be explained by the odor responses of forebrain neurons. These findings suggest that odor stimulation not only alters the dynamics of the forebrain circuitry but also changes the way habenula integrate its forebrain inputs, and further modulates the state of ongoing habenular activity.

## DISCUSSION

In this study, we revealed the forebrain circuitry underlying ongoing habenular activity and showed that this inter-connected network can generate structured ongoing activity with strong synchrony and anti-synchrony that is stable over time. Our results further showed that chemosensory stimuli can modulate brain activity not only by activating the olfactory pathway, but also by suppressing the ongoing activity of habenular networks and its input regions in the forebrain that are ancestral to mammalian limbic system. Interestingly, our micro-stimulation experiments also revealed that the specific communication between habenula and its forebrain inputs is dampened in the presence of odor stimulation. Disruption of ongoing (or resting state) brain activity with sensory stimuli has also been observed in human subjects [3, 6, 7], suggesting a high-level of evolutionary conservation (or convergent evolution) of vertebrate brain dynamics and the way they interact with the sensory world. Such interactions could serve multiple purposes. First, disruption of ongoing or resting brain state with a salient sensory stimulus, can facilitate state transitions, when an animal needs to quickly attend, adapt and respond to salient environmental changes. In fact, such a role of the habenula in behavioral flexibility and in adapting to new conditions has been demonstrated in both zebrafish [8, 64, 65, 72, 73, 76] and mammals [1, 134, 135]. Second, dampening of ongoing habenular and forebrain activity might increase the sensitivity of these networks to incoming sensory information by reducing background noise levels. This would, in turn, increase the robustness of sensory representations to incoming stimuli following first encounter, as it is often the case with olfactory stimuli that are received by repeated exposure to odor plumes [136–138]. Finally, resetting the ongoing activity of limbic forebrain networks, might be an important mechanism allowing animals to make new associations, when conditions are altered, as it has been shown during reversal learning [1, 75, 78, 135]. How does the ongoing activity of these networks adapt during behavioral plasticity and learning will require well-established behavioral assays and simultaneous imaging of forebrain and habenular activity in juvenile zebrafish, that can perform cognitively demanding learning tasks [75, 95–97]. Moreover, given the multi-sensory responses of habenular circuits [52], it will also be interesting to test whether such interactions between sensory stimuli and ongoing brain activity generalize to other sensory modalities (e.g. vision, auditory), in future studies.

Anatomical projections from the olfactory bulb [38, 111], entopeduncular nucleus [41, 49, 55, 58], and lateral hypothalamus [41, 47, 100] to habenula have been shown by several past studies. In addition to these anatomically confirmed regions, our results showed that Dl and Dm, zebrafish homologs of mammalian hippocampus and amygdala, are the main dorsal forebrain regions that are strongly correlated with ongoing habenular activity. We also confirmed these functional connectivity measures based on correlations, by further micro-stimulation of Dl and Dm that activate up to ~30% of habenular neurons. It is yet to be discovered whether distinct habenular sub-circuits encode information from different forebrain regions. To our knowledge no direct anatomical projections from Dl or Dm were reported to innervate habenula in zebrafish. Nevertheless, Dm neurons were shown to innervate entopeduncular nucleus and hypothalamus [106] both of which were shown to project to habenula in zebrafish and mammals [41, 47, 56, 58, 100, 102, 125, 128]. Dl neurons are a larger group of neurons, which are likely functionally heterogenous due to large size of Dl region. All studies of Dl in teleost fish highlight a homology of this structure with hippocampal-entorhinal circuitry related to episodic memory and spatial navigation [119, 120, 123, 126, 139]. In fact, similar to our findings in zebrafish, functional coupling between hippocampal circuitry and habenula has been shown also in rodents [140–142]. All these results suggest that there might be direct or indirect connections from hippocampal circuitry to habenula. Extensive anatomical reconstructions of Dl neurons, by using newly generated electron- and light-microscopy imaging data sets from zebrafish brains [105, 143], will shed more light onto direct or multi-synaptic anatomical connections between the habenula and the hippocampal system. This may also highlight new avenues of research on the role of habenula for episodic memories and spatial navigation.

As a small vertebrate, zebrafish exhibit general principles of vertebrate forebrain architecture. Most studies of zebrafish forebrain circuitry have investigated adult zebrafish with sophisticated behaviors and large brains that are challenging to access using optical imaging methods and requires surgery [125, 144, 145]. The forebrain, especially the dorsal telencephalon, of transparent 5-10 days old zebrafish larvae is not yet fully developed [52, 128, 146]. Hence, using 3 to 4-week-old juvenile zebrafish provides an alternative approach to study a relatively developed forebrain network [52, 128, 146] that is still optically accessible to imaging the entire forebrain, including habenula, as we demonstrated in this study. This approach allowed us to identify functionally distinct populations of forebrain neurons, which also overlapped with distinct forebrain regions that we could identify based on anatomical landmarks. Our calcium imaging results allowed us to relate some of these functionally defined zones in juvenile zebrafish forebrain with adult zebrafish, other teleosts and even mammalian forebrain structures. Given the ability of juvenile zebrafish to perform cognitively demanding behaviors such as social interactions [97, 98] and learning [75, 95, 96], while allowing non-invasive functional imaging, we hope that future studies will shed further light onto the function and the connectivity of forebrain networks during these behaviors.

Altogether our results highlight the activity and connectivity of forebrain networks underlying ongoing habenular activity and revealed how habenula integrates olfactory and limbic information. Our findings further support the idea that the habenula serve as a main hub [44, 74], which can integrate sensory and non-sensory information from diverse forebrain regions and relays this integrated information to its downstream targets regulating animal behavior [41, 43–48, 50, 58]. Given the important role of habenula in adaptive behaviours and mood disorders [147–150], our results suggest that sensory experience might provide a non-invasive pathway to modulate habenular activity, perhaps also in humans. Future studies with high-throughput and high-resolution brain imaging methods in mammals, may answer whether sensory information can perturb the dynamics of ongoing activity of mammalian habenula.

## ACKNOWLEDGEMENTS

We thank M. Ahrens (HHMI, Janelia Farm, USA), K. Kawakami (National Institute of Genetics, Japan) for transgenic lines. We thank S. Eggen, M. Andresen, V. Nguyen and our fish facility support team for technical assistance. We thank the Yaksi lab members for stimulating discussions. We thank Steffen Kandler and Nathalie Jurisch-Yaksi for their feedback on the manuscript. This work was funded by ERC starting grant 335561 (E.Y.), Helse Midt-Norge Samarbeidsorganet grant (E.Y.), RCN FRIPRO Research Grant 239973 (E.Y.), Boehringer Ingelheim Fonds (S.K.J.). Work in the E.Y. lab is funded by the Kavli Institute for Systems Neuroscience at NTNU.

## AUTHOR CONTRIBUTIONS

Conceptualization, E.M.B., S.K.J., E.Y.; Methodology and data, E.M.B., S.K.J., E.Y.; Data Analysis, E.M.B., S.K.J., K.T.P.C., E.Y.; Investigation, all authors; Writing, E.M.B., E.Y.; Review & Editing, all authors; Funding Acquisition and Supervision, E.Y.

## MATERIALS AND METHODS

### Fish husbandry

NFSA (Norwegian Food Safety Authority) has approved the animal facility and fish maintenance. Fish were kept in 3,5L tanks in a Tecniplast ZebTec Multilinking System. Constant conditions were maintained: 28.5°C, pH 7.2, 700μSiemens. 14:10 hour light/dark cycle was preserved. Dry food (SDS100 up to 14dpf and SDS 400 for adult animals, Tecnilab BMI, the Netherlands) was given to fish twice a day, in addition to Artemia nauplii (Grade 0, Platinum Label, Argent Laboratories, Redmond, USA) once a day. From fertilization to 3dpf (days post fertilization) larvae were kept in a Petri dish with egg water (1.2g marine salt in 20L RO water, 1:1000 0.1% methylene blue) and between 3 and 5dpf in artificial fish water (AFW: 1.2 g marine salt in 20L RO water). Juvenile (3 to 4-week-old) zebrafish were used for the experiments.

For calcium imaging, Tg(elavl3:GCaMP5) (Jetti, Vendrell-Llopis, and Yaksi 2014) and Tg(elavl3:GCaMP6s) (Park et al. 2000) zebrafish lines were used.

### Two-photon calcium imaging

For in-vivo imaging, fish were embedded in 2-2,5% low-melting-point agarose (LMP, Fisher Scientific) in a recording chamber (Fluorodish, World Precision Instruments). To ensure odor to arrive to the nostrils, the LMP agarose was removed carefully in front of the nose, after solidifying for 20min. The constant perfusion of AFW bubbled with carbogen (95% O2 and 5%CO2) was maintained during the experiment.

For in-vitro imaging, brain explants were prepared as described in section “Dorsal dissection of juvenile zebrafish brain” below. The preparation is constantly perfused with artificial cerebrospinal fluid (ACSF) AFW was bubbled with carbogen (95% O2 and 5%CO2) throughout the experiment.

Two-photon microscopes were used for calcium imaging: Scientifica Inc, with a 16x water immersion objective; Nikon, NA 0.8, LWD 3.0 and Zeiss 7 MP with a 20x water immersion objective W Plan-Apochromat, NA 1.0. For excitation a mode-locked Ti:Sapphire laser (MaiTai Spectra-Physics) was tuned to 920nm. Recordings were performed as either single plane or volumetric recording (6-8 planes with a Piezo (Physik Instrumente (PI)). The acquisition rate was 6,4Hz for single plane recordings (image size 256×512 pixels) and between 2.3-3.4Hz per plane for volumetric scans (average image size 1536×750).

### Odor preparation

Our odor panel consisted of food odor, skin extract, urea, bile-acid mixture, amino acid mixture, and ammonium chloride. All odorants were purchased from Sigma Aldrich. Apart from food odor and skin extract, all odors were used at 10-4M concentration. Food odor was prepared using commercially available fish food; 1g of food particles was incubated in 50ml of fish water (FW) for at least 1 hour, filtered through filter paper, and diluted to 1:50. For skin extract (Kermen et al. 2020), adult zebrafish were first euthanized in ice-cold water and decapitated, the skin was peeled off from the body. 1g of skin was incubated in 2ml of AFW and was vortexed at 1300rpm for 1h at 4°C. After, the skin extract was dissolved in 50ml of AFW and filtered through the filter paper. All odors were prepared from the frozen stocks immediately before use.

### Odor delivery

The stimulation tube was positioned in front of the nose and the stimulus was delivered for 30s. The stimulation was performed with HPLC injection valve (Valco Instruments) controlled with Arduino Due. Before each experiment, a trial with fluorescein (10-4M in AFW) was performed to determine precise onset of odor delivery.

### Micro-electrode stimulation

Electrodes were pulled from a borosilicate glass microcapillary (1.00 mm; World Precision Instruments) using a laser puller (Sutter Instruments co. Model P-2000). Electrodes tips were were broken so that the final tip diameter of the stimulating electrode was around 10-15μm. The electrode was filled with artificial cerebro-spinal fluid (ACSF). Two-separate electrodes were connected to two poles of the ISO-flex stimulus isolator (A.M.P.I.), and electrodes were glued by two-component epoxy so that the electrode tips are close to each other (within 1-2 mm). Only a single electrode tip was inserted and positioned in the brain explant using a micromanipulator (Scientifica, UK), and the other tip was left outside the brain. A train of short current pulses (2-5μA) for 50 milliseconds duration and 20 seconds inter stimulus intervals was applied to the target brain region, using a computer-controlled Arduino Due as a stimulus generator.

### Dissection of juvenile zebrafish brain explant

All animals (3-4 weeks post fertilization) were anesthetized in ice-cold artificial fish water (AFW) and euthanized by decapitation in oxygenated artificial cerebro-spinal fluid (ACSF). The dissection was carried out in ACSF. After removing jaws and eyes, muscle tissue, gills and fat were cleaned to allow oxygen diffusions to the brain explant. Next, the skin tissue, bone and dura mater covering the forebrain was removed from the dorsal side, allowing electrode access.

### Data analysis

Two-photon microscopy images were aligned using a method described in (Reiten et al. 2017; Verdugo et al. 2019). Occasional XY drift was corrected, based on “hierarchical model-based motion estimation” (Bergen et al. 1992). Recordings were then visually inspected for remaining motion and Z-drift, recordings with remaining motion artifacts were discarded. Regions of interest (ROIs) corresponding to neurons were automatically detected using a template matching algorithm (Jetti, Vendrell-Llopis, and Yaksi 2014; Verdugo et al. 2019), and visually confirmed. To calculate the time course of each neuron, pixels belonging to each ROI were averaged over time. For each ROI, fractional change in fluorescence (ΔF/F) relative to baseline was calculated.

Neurons were clustered into functional clusters using k-means clustering algorithm in MATLAB (Figure 1A, Figure 2B, Figure 3E) (Jetti, Vendrell-Llopis, and Yaksi 2014). For Figure 2 an average activity of all neurons in each cluster of the habenula was averaged. This population vector was used as a reference activity and correlated with each neuron in the telencephalon.

Delineation of brain regions in the telencephalon was done manually on the raw image of the brain (as in Figure 2E) based on anatomical landmarks (Rupp, Wullimann, and Reichert 1996) described in previous studies in zebrafish and other teleost fish (Diaz-Verdugo et al. 2019; Huang et al. 2020; Wullimann and Mueller 2004; Elliott et al. 2017; Fotowat et al. 2019; Rodriguez-Exposito et al. 2017).

Cluster selectivity was calculated to quantify the overlap of manually delineated brain regions with functional clusters of neurons identified using k-means clustering based on their spontaneous activity. Cluster selectivity index is the life-time sparseness (Jetti, Vendrell-Llopis, and Yaksi 2014) for the distribution of neurons within an anatomically defined brain region across the k-means clusters. If cluster selectivity is 1, it means that neurons of an anatomically identified brain region belong to one functional cluster. If cluster selectivity is 0, all neurons of an anatomically identified brain region are equally distributed into all functional k-means clusters.

Cluster fidelity was calculated by measuring the probability of pairs of neurons being in the same cluster during two different time periods (Jetti, Vendrell-Llopis, and Yaksi 2014). We compared the cluster fidelity of real k-means clusters with shuffled the cluster identities of same neurons.

In Figure 4, neurons were stimulated 6 times with each stimulus: odor, micro-stimulation and odor + micro-stimulation. Neurons were considered as responsive if their response (5s) exceeded 2 standard deviations above the 5s-baseline period preceding the stimulation. To calculate how odor and micro-stimulation influenced each other, we compared whether these two different stimuli can significantly (using Wilcoxon signed rank test) change each other when delivered simultaneously (odor + micro-stimulation).

For Figure 5, odor responses of habenular neurons were classified by correlating their odor response with a step function, as previously described in (Cheng et al. 2017). Top 5% of habenular neurons with highest or lowest correlations were considered as strongly excited or inhibited, respectively. Averaged ongoing activity preceding the stimulation in those classified habenular neurons was used as a reference vector and correlated with each forebrain neuron to identify forebrain neurons that are highly correlated with excited or inhibited habenular neurons.

### Statistics

Statistical analysis was done using MATLAB; p-values are represented in the figure legends as (*p<0.05, **p<0.01, ***p<0.001). Wilcoxon ranksum test was used for non-paired comparisons and Wilcoxon signed rank test for paired comparisons.

All analysis was performed with Fiji and MATLAB.

