## Supplemental Figures for "Ongoing habenular activity is driven by forebrain networks and modulated by olfactory stimuli"

### Supplementary Figures

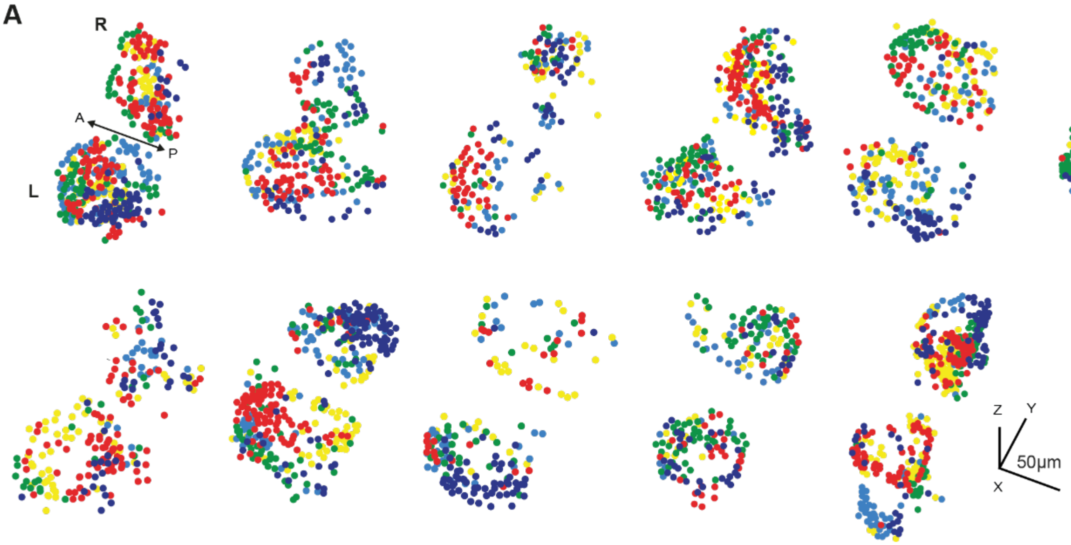


**Figure S1. Functional clusters of habenular neurons in all imaged zebrafish.**

(A) Three-dimensional reconstruction of functional clusters of habenular neurons identified using k-means clustering. n=11 different animals. Neurons similar ongoing activity are color-coded based on their functional cluster identity. L-left; R-right hemisphere, A-anterior, P-posterior.

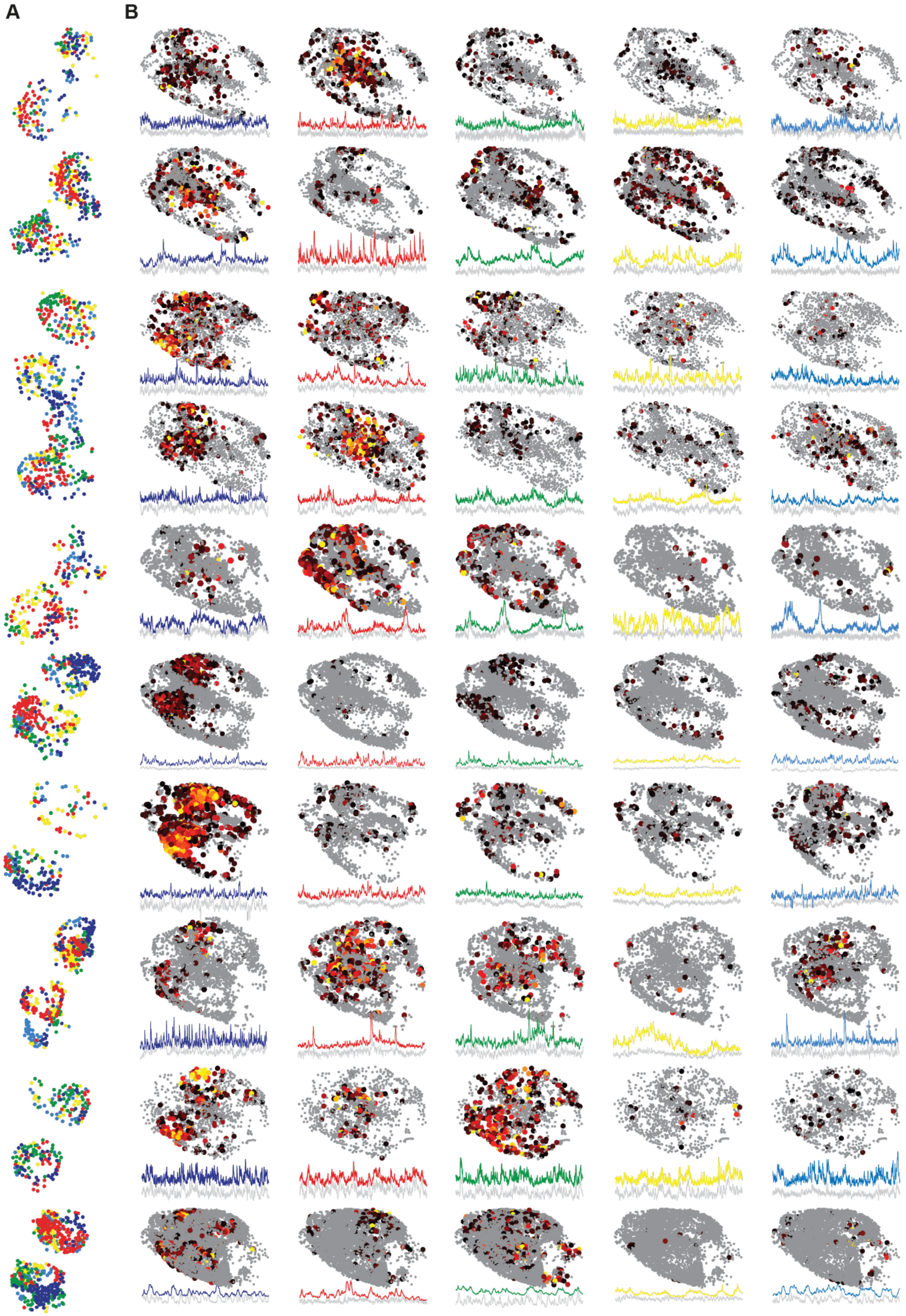


**Figure S2. Forebrain neurons that are correlated with the ongoing activity of functional habenular clusters, in all imaged zebrafish**

(A) Three-dimensional reconstruction of habenular neurons detected in *Tg(elavl3:GCaMP6s)* zebrafish line, clustered with k-means clustering. Colors represent neural clusters with similar ongoing activity. L-left; R-right hemisphere. 315 ± 35 (mean ± SEM) habenular neurons were imaged in each fish (n=11 fish).

(B) Three-dimensional reconstruction of forebrain neurons that are strongly correlated (Pearson's correlation >0.1) to average ongoing activity of different habenular clusters in B. Warm colors represent stronger correlations. 2135 ± 345 (mean ± SEM) forebrain neurons were imaged in each fish (n=11 fish). Color-coded traces represent the average activity of forebrain neurons in each cluster. Grey traces represent the average activity of corresponding habenular clusters in A. Note that the ongoing activity of identified forebrain neurons and habenular clusters are highly similar.

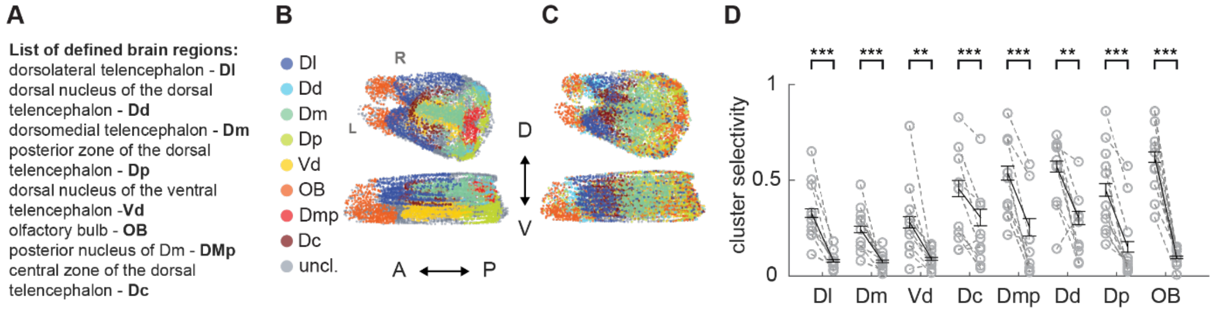


**Figure S3. Forebrain regions identified by using anatomical landmarks correspond to functional clusters of forebrain neurons identified by k-means clustering.**

(A) A list of anatomically identified zebrafish forebrain regions and their abbreviations.

(B) Three-dimensional reconstruction of forebrain neurons identified by using anatomical landmarks. Top – dorsal view, bottom – coronal view. L-left; R-right hemisphere. A – anterior, P – posterior, D- dorsal, V-ventral. Colors corresponds to individual forebrain regions in A.

(C) Three-dimensional reconstruction of forebrain neurons that are clustered by k-means functional clustering (with n=8 clusters) of their ongoing neural activity. Each arbitrary color represents functional forebrain cluster.

(D) Neurons of anatomically identified forebrain regions exhibit cluster selectivity that are significantly higher than chance levels.

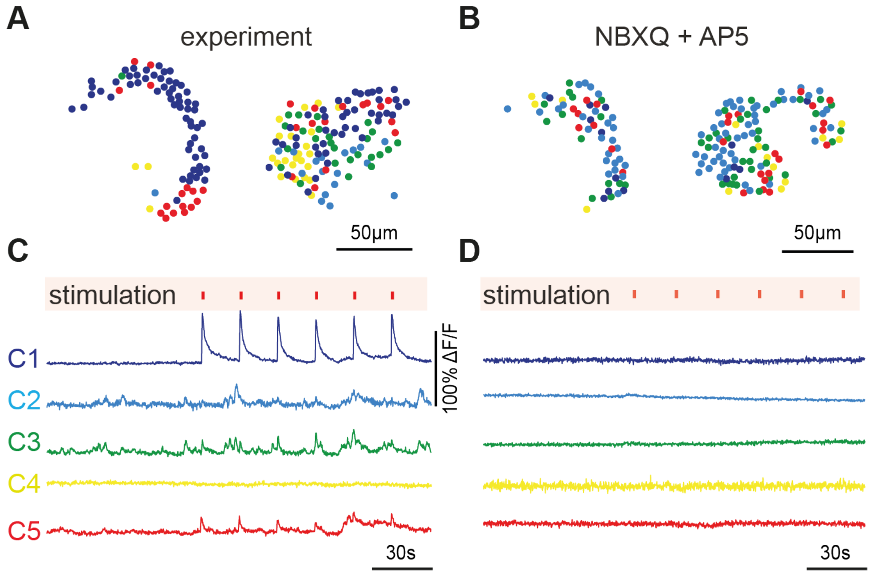


**Figure S4. Responses of habenular neurons upon forebrain micro-stimulation is eliminated by blockers of glutamatergic neurotransmission NBXQ + AP5.**

(A) Representative example of habenular neurons detected in *Tg(eval3:GCaMP5)* zebrafish line during forebrain micro-stimulation, clustered with k-means clustering. Neurons are color-coded based on their cluster identity (C1-5).

(B) Average activity of each habenular functional cluster (colors corresponds to panel A) during 6 consecutive, 50ms forebrain micro-stimulations. Clusters are defined by k-means clustering. Forebrain micro-stimulations are marked in red.

(C) Representative example of habenular neurons detected in *Tg(eval3:GCaMP5)* zebrafish line during forebrain micro-stimulation, in the presence of glutamatergic blockers NBXQ(5µm) + AP5(25µm ).

(D) Average activity of each habenular functional cluster (colors corresponds to panel C) during 6 consecutive, 50ms forebrain micro-stimulations. Note that in the presence of NBXQ(5µm) + AP5(25µm ), no habenular responses are detected.

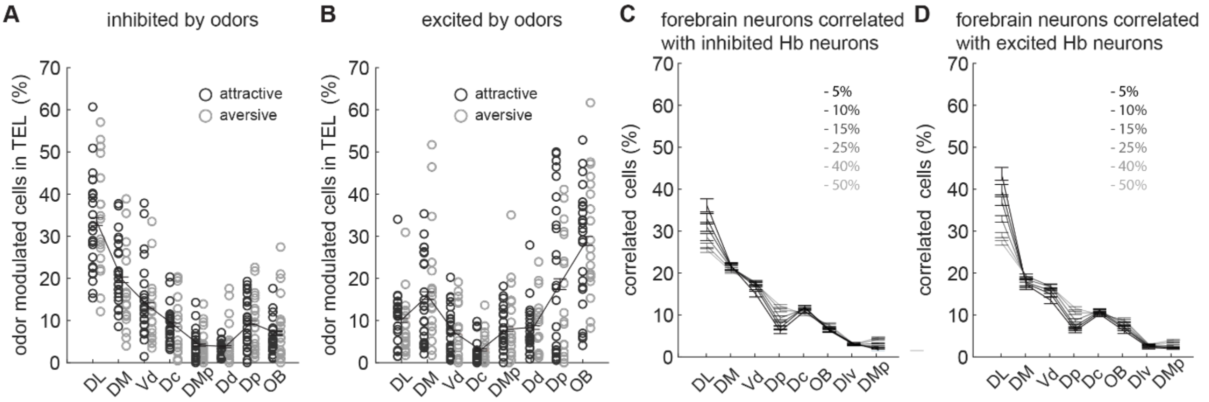


**Figure S5. Distribution of forebrain neurons into anatomically identified forebrain regions, based on their odor responses, or their synchrony with odor modulated habenular neurons.**

(A) Anatomical distribution of forebrain neurons that are most (top 5%) inhibited by odors in 10 fish. Black and gray circles represent attractive and aversive odors, respectively. Black lines represent mean ± SEM.

(B) Anatomical distribution of forebrain neurons that are most (top 5%) excited by odors in 10 fish. Black and gray circles represent attractive and aversive odors, respectively. Black lines represent mean ± SEM.

(C) Anatomical distribution of forebrain neurons with varying correlations (between 5 and 50%) to ongoing activity of habenular neurons that are inhibited by odors in n=10 fish. Relates to Figure 5E.

(D) Anatomical distribution of forebrain neurons with varying correlations (between 5 and 50%) to ongoing activity of habenular neurons that are excited by odors in n=10 fish. Relates to Figure 5E. Dl – dorsolateral telencephalon, Dd – dorsal nucleus of the dorsal telencephalon, Dm – dorsomedial telencephalon, Dp – posterior zone of the dorsal telencephalon, Vd – dorsal nucleus of the ventral telencephalon, OB – olfactory bulb, Dmp – posterior nucleus of dorsomedial telencephalon, Dc – central zone of the dorsal telencephalon.
